# Evidence that the *Kuenenia stuttgartiensis* encapsulin does not protect against NO damage

**DOI:** 10.64898/2026.06.22.733830

**Authors:** John Tracey, Tobias W. Giessen, Bess B. Ward

**Affiliations:** Princeton University, Department of Geosciences, Guyot Hall, Princeton, NJ 08544, USA; University of Chicago, Department of the Geophysical Sciences, Chicago, IL, 60637, USA; University of Chicago, Climate Systems Engineering initiative, Chicago, IL, 60637, USA; University of Chicago, Institute of Climate and Sustainable Growth, Chicago, IL, 60637, USA; Harvard Medical School, Department of Systems Biology, Boston, MA 02115, USA; Wyss Institute for Biologically Inspired Engineering, Boston, MA, 02115, USA; Department of Biological Chemistry, University of Michigan, Ann Arbor, MI, 48109, USA

**Keywords:** Anammox, Encapsulin, Nitric Oxide, Compartmentalization

## Abstract

A paradigm shift is underway in microbiology: many prokaryotes, long considered to lack the compartmentalization present in all eukaryotic life, have been found to possess a great diversity of protein based intracellular compartments. Notably, the genomes of many marine and freshwater anaerobic ammonium oxidizing (anammox) bacteria encode one of these compartmentalization strategies; encapsulin nanocompartments. These systems’ structure suggests a role for anammox encapsulins in the anammox metabolism, a process of global biogeochemical significance, which results in the loss of biologically available nitrogen from aquatic environments. Here we test if the most common anammox encapsulin architecture could provide a mechanism to detoxify NO, one of the reactive intermediates produced in the core anammox metabolism. Through experiments in which the *Kuenenia stuttgartiensis* encapsulin was heterologously expressed by an inducible plasmid in *E. coli*, we show evidence that suggests the *K. stuttgartiensis* encapsulin provides no protection from NO.

## Introduction

Anammox bacteria perform one of only two known biological pathways that return fixed (biologically available) nitrogen back to largely unusable N_2_. As a result, anammox bacteria play a large role in determining the fixed N budget in many environments. The core metabolism involves three redox half-reactions that in combination oxidize ammonium with nitrite to create N_2_ gas^1^. These steps, listed in order with their respective enzymes, are:

1. NO_2_^-^ + 2H^+^ + e^-^ = NO + H_2_O (Nitrite reductase, NIR)
2. NO + NH_4_^+^ + 2H^+^ + 3e^-^ = N_2_H_4_ + H_2_O (Hydrazine synthase, HZS)
3. N_2_H_4_ = N_2_ + 4H^+^ + 4e^-^ (Hydrazine dehydrogenase, HDH, HAO-like)
4. NO_2_^-^ + NH_4_^+^ = N_2_ + 2H_2_O (Net)

The core anammox metabolism involves two extremely reactive compounds, hydrazine (N_2_H_4_), which has a propensity to react with phosphate to produce free radicals^2^, and the reactive nitrogen species NO. Due to the danger these compounds pose to biological materials, these compounds’ toxicity was originally proposed as the rationale for the anammoxosome, a membrane-bound structure which, as observed via electron microscopy, occupies most of the cell volume of anammox bacteria^3^. The anammoxosome membrane is composed of ladderane lipids, which possess an unusual planar ring structure that folds into a series of square planar steps resembling a ladder. This unique membrane structure was thought to make the anammoxosome membrane more impermeable to free radical escape, since the square kinks of adjacent ladders can be more densely packed than standard fatty acids chains^3,4^. However, more recent studies have shown that ladderane lipids likely function to maintain a pH gradient across the anammoxosome membrane, not protect against reactive compound escape^5^.

Prokaryotes, with a few exceptions such as the anammoxosome, were long considered incapable of the compartmentalization achieved by eukaryotic membrane-bound organelles. However, it is becoming increasingly clear that a great diversity of prokaryotes achieve a high degree of subcellular organization and compartmentalization^6,7^. Of the prokaryotes with compartmentalization strategies, most have been found to employ protein-based, rather than lipid-based, compartments, which fall into two main classes: bacterial microcompartments (BMCs) and encapsulin nanocompartments^8^. These compartments give prokaryotes many of the same benefits of eukaryotic organelles such as an improved ability to regulate metabolic reactions due to a concentrated microenvironment containing enzymes and reactants, the capacity to store enzymes or reactants for long periods of time, greater efficiency in transporting reactants and enzyme regulators since they can be directed to one location, and the power to synchronously perform incompatible reactions in separate differentiated reaction microenvironments^9,10^.

Encapsulin nanocompartments are hollow protein shells with an icosahedral geometry^9^. Over 6,000 encapsulins have been bioinformatically discovered so far^11^. These proteins are further classified into four families based on distinctions in encapsulin amino acid sequence, geometry, and operon architecture. The members of these families span 31 bacterial and 4 archaeal phyla, have triangulation numbers of T=1, T=3, and T=4, share the HK97 (Hong Kong 97) phage-like fold previously thought to be unique to viruses and bacteriophages^11^, and range in diameter from 24 to 42 nm^8^. Importantly, nearly all encapsulins enclose one or more specific cargo enzymes, which are normally targeted to the encapsulin complex interior by short, highly conserved targeting peptides at the C- or N-terminus of the cargo proteins or by longer N-terminal cargo-loading domains^12–15^. These cargo enzymes are usually encoded immediately before or after the encapsulin monomer.

Of the thousands of encapsulin protein sequences currently known, eight unique sequences have been identified in anaerobic ammonium oxidizing (anammox) bacteria^9,11,16^ (See supplementary material for an explanation of this tally of anammox encapsulins). Five of the anammox encapsulin systems currently known possess an unusual double fusion protein architecture^9,11^ (Figure 1). These systems encode an encapsulin/cytochrome c fusion protein, containing two CXXCH heme-binding sites, and a nitrite reductase-like (NIR)/HydroxylAmine Oxidoreductase (HAO)-like fusion protein encoded immediately up or downstream of the encapsulin gene. All five of the anammox encapsulins of this type belong to freshwater anammox candidate species, while the other anammox encapsulins, including two from the marine genus *Scalindua*, contain variations on this theme. For example, *Scalindua rubra* encodes an encapsulin without a cytochrome c domain, immediately followed by two nitrite reductase-like enzymes, while *Scalindua* sp. SCAELEC01 167 (also known as sp. AMX11) encodes a NIR/HAO fusion enzyme, an encapsulin fused to a hypothetical protein domain, and an additional nitrite reductase^16^. Two pieces of evidence suggest that anammox encapsulins are located in the anammoxosome. First, a proteomics study performed on an anammoxosome enrichment has demonstrated the presence of encapsulins within the anammoxosome proteome, albeit at low levels^17^. In addition, small ∼25 nm diameter electron and iron-rich regions have been observed throughout the anammoxosome^18^. These regions could be encapsulins, based on the approximate size agreement with electron microscopy data of the purified *Kuenenia stuttgartiensis* encapsulin and the prevalence of heme groups discovered in many of the known anammox encapsulins^9,16^.

**Figure 1:**
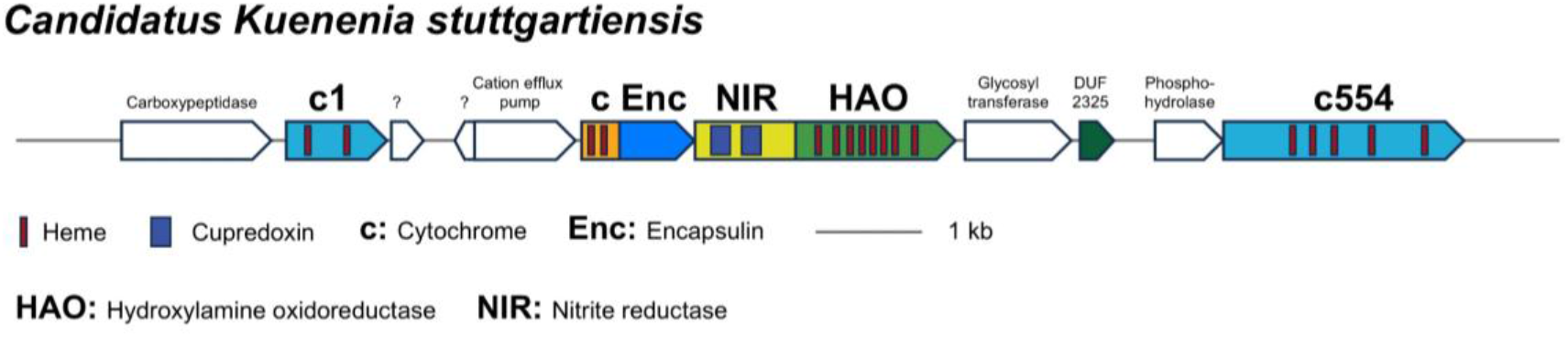
The *Kuenenia stuttgartiensis* encapsulin system, one of five known anammox encapsulins that include both encapsulin/cytochrome and putative NIR/HAO fusion cargo proteins. c1 and c554 are putative secondary cargo identified in a previous publication^9^. The protein sequence of cEnc is included in the supplementary materials.

Of the known anammox encapsulin systems, the *Kuenenia stuttgartiensis* system is the best studied. The *K. stuttgartiensis* encapsulin system consists of a cytochrome-encapsulin fusion protein (cEnc) with CXXCH binding sites followed by a putative NIR/HAO fusion protein that is thought to be the encapsulin cargo. Based on a SignalP search, cEnc also contains an N terminal Sec-type signaling sequence^19^. The Sec pathway is one of the primary peptide secretion pathways in bacteria^20^. Sec-peptide carrying proteins are usually transported across a membrane to their destination in an unfolded state^21^. The presence of a Sec peptide therefore implies that *K. stuttgartiensis* encapsulin shell proteins are transported unfolded across a membrane, likely the anammoxosome membrane. Previous studies have heterologously expressed the *K. stuttgartiensis* cEnc and NIR/HAO and purified cEnc and the NIR domain of the NIR/HAO fusion from *E. coli* cultures^9^. Purified cEnc was also observed to self-assemble into 33-nm nanocompartments by transmission electron microscopy (TEM) and successfully incorporate heme as shown by observed dithionite reduction in UV-spectroscopy^9^.

The likely co-localization of anammox encapsulins with the anammoxosome, the proposed NIR/HAO cargo protein, and the presence of toxic intermediates like NO and N_2_H_4_ in the core anammox metabolism led us to test our hypothesis for an interconnection between anammox encapsulins, the core anammox metabolism, and detoxification. Since NO is an intermediate in central anammox metabolism it is reasonable to suspect that anammox bacteria would have a detoxification system if NO concentration exceeded that required by HZS.

## Methods

### Strain development

Our hypothesis was investigated by expressing the *Kuenenia stuttgartiensis* cEnc protein on an inducible plasmid in *E. coli*. This strategy was chosen because anammox cultures have such low doubling times (7-20 days)^22^ that even with extensive culture hardware engineering can only reach 2 - 4 days^23^. *E. coli* offers doubling times on the order of minutes to hours and allows IPTG inducible gene expression^24^. In addition to testing our NO protection hypothesis using the heterologously expressed *Kuenenia* cEnc, we also developed strains that included the NIR/HAO putative cargo protein. All strains are listed below (Table 1) and were created following a previous publication^9^. In brief, Gibson Assembly was used to build constructs joining encapsulin components and the plasmid vector. All gBlock encapsulin and putative cargo-bearing gene fragments were joined with NdeI and PacI digested pETDuet1 vectors. Chemically competent *E. coli* Turbo cells were then transformed with the vectors and gene insertion was confirmed by sequencing. Validated plasmids were used to transform *E. coli* BL21 (DE3) Star cells (∼0.5 ng total DNA) to create the strains listed in Table 1. Since pETDuet1 encodes ampicillin resistance, successful transformation was confirmed by successful growth in LB media with ampicillin (100 µg ml^-1^). Successful expression and purification of cEnc and the NIR domain was confirmed via gel electrophoresis^9^. Successful encapsulin formation was confirmed via TEM imaging^9^. After strain construction, stocks were created by freezing 1 mL of dense culture (OD > 0.5) in glyercol and storing at -80°C. Strains TG226 and TG227 contain the full-length *K. stuttgartiensis* cytochrome/encapsulin fusion which includes a Sec signaling peptide. This Sec signaling peptide was removed in strains TG228 and TG229. While all strains could be successfully grown with IPTG induction we focused our NO protection hypothesis testing on strain TG228 because that was the only strain where successful purification, assembly, and heme incorporation had been shown previously^9^ and the decline in growth rate observed in signal peptide expressing strains TG226 and TG227 (Figure 2) could be a confounding factor.

**Table 1:** Description of Anammox Constructs

| Strain ID | Insert | Description |
| --- | --- | --- |
| TG226 | Ks-cEnc | *K. stuttgartiensis* cytochrome/encapsulin fusion with Sec signaling peptide |
| TG227 | Ks-cEnc-NIR/HAO | *K. stuttgartiensis* cytochrome/encapsulin fusion and NIR/HAO cargo, with Sec signaling peptide |
| TG228 | Ks-cEnc-no signal | *K. stuttgartiensis* cytochrome/encapsulin fusion, no signaling peptide |
| TG229 | Ks-cEnc-NIR/HAO-no signal | *K. stuttgartiensis* cytochrome/encapsulin fusion and NIR/HAO cargo, no signaling peptide |
| TG306 | - | Wild Type (No plasmid) |

**Figure 2:**
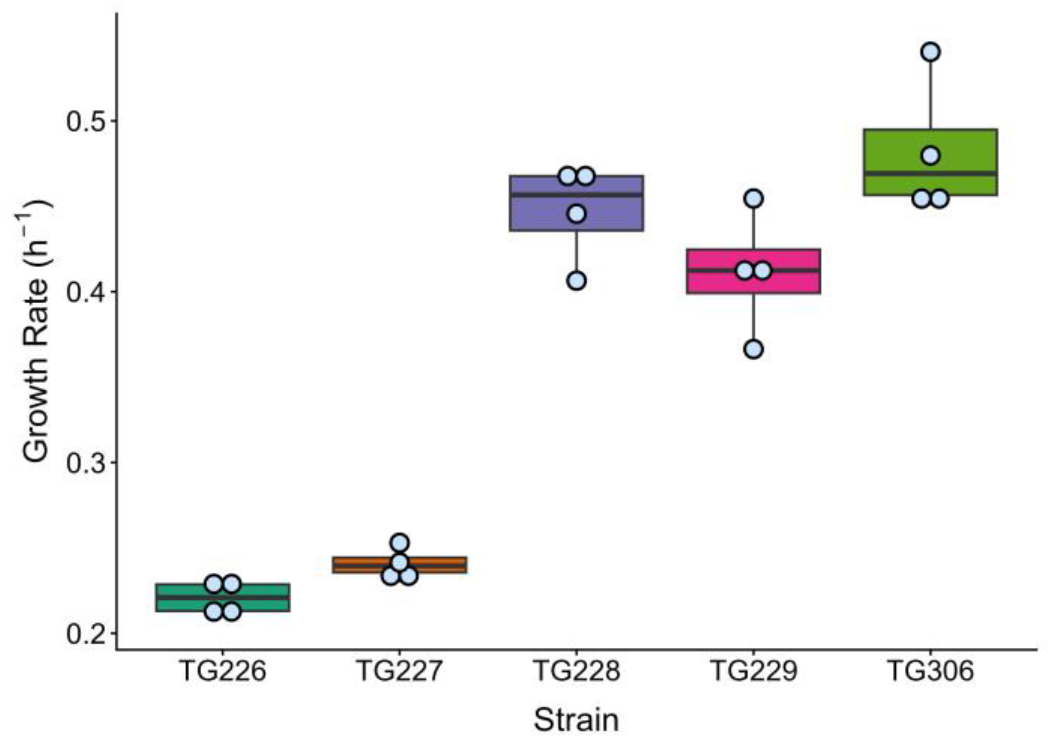
Expression of Sec signaling domain in strains TG226 and TG227 lowers growth rate. Each point represents the growth rate calculated from one plate of a 12-well plate. Whiskers show minimum and maximum growth rates, boxes show the middle 50% of the growth rate data, while solid black lines show the median growth rate for each strain.

### Growth Experiments Investigating Growth Curve Reproducibility and Sec Peptide

Strains TG226, TG227, TG228, TG229, and TG306 were inoculated from frozen stocks into 10 mL aerobic M9 medium with added ampicillin (100 µg ml^-1^) for strains TG226, TG227, TG228, and TG229. TG 306 received no ampicillin. Cultures were grown overnight at 37°C and 50 rpm. The following day, overnight cultures were inoculated 1:50 into M9 (+/- ampicillin as before) across each well (3 mL M9 per well) of a 12 well tissue culture plate (Corning™ Costar™). The M9 recipe is included in the supplement. Expression was induced by adding IPTG to 0.1 mM to all strains. All strains were grown in duplicate. The plate was then grown at 30°C in a Biotek Synergy HTX plate reader for 24 hours with OD 660 measurements taken every 10 minutes, with no path length correction, and shaking at 180 rpm. This plate reader experiment was repeated once giving a total of four samples per strain. Growth rates were calculated by first identifying the period of exponential growth in each well’s growth curve (i.e. the part of the ln(OD 660) vs. time curve that is linear) and then computing a linear regression between the natural logarithm of the OD 660 and time over that time range.

### NOC - 18 Addition Experiments: Testing for Protection from NO

M9 medium was used for NOC-18 addition experiments so that the nitrogen composition of the medium was defined. Deoxygenated M9 medium with ampicillin was prepared by microwaving 500 mL of previously autoclaved medium for nine minutes on a Panasonic Sensor 1250 microwave set at Power level 10. This medium was then sonicated for 2 minutes under vacuum on a Branson 1200 sonicator and finally bubbled for 30 minutes with nitrogen gas. After bubbling, the medium was immediately placed into a Coy Laboratories glove box under an atmosphere of N_2_. Within the glove box, the medium was filtered and aliquoted into sealed and crimped Hungate tubes.

The *E. coli* cultures of strains TG 228 and TG 306 were gradually acclimated to anaerobic conditions. Aerobic overnight starter cultures were grown overnight at 37°C on LB / Agar plates supplemented with ampicillin at 100 µg ml^-1^. A single colony from this plate was then inoculated into 15 mL of aerobic M9 medium and grown at 37°C, 180 rpm until the OD600 reached 0.3. This culture was then transferred at a 1:50 ratio into sealed Hungate tubes containing 10 mL anaerobic M9 and grown at 37°C overnight. The following day, overnight anaerobic cultures were inoculated 1:10 into 10 mL of anaerobic M9 and grown at 30°C. Expression was then induced by adding IPTG to 0.1 mM and hemin and FeSO_4_ cofactors were added to 2 µM. NOC - 18 was weighed as a powder and placed in the glove box overnight to degas. Immediately prior to addition, NOC - 18 was dissolved in 1 µM NaOH, sealed in a Hungate tube, and added to the test strains at 1 mM final concentration at 5 hours. OD 600 values were measured throughout a ten-hour experiment on a Genesys 30 spectrophotometer (Thermo-Fisher). Aerobic experiments were performed as above except for the use of aerobic M9 medium and different concentrations of NOC-18.

## Results

### Engineered *E. coli* Strains Grow Reproducibly While Sec Signaling Domain is Associated with Growth Rate Differences Across Strains

Before conducting experiments to directly test our hypotheses about the role of anammox encapsulins in cellular detoxification and anammox central metabolism, we performed growth experiments to check that the engineered *E. coli* strains had reproducible growth rates after induction with IPTG. All strains (Table 1) had highly reproducible growth rates (Figure 2). Interestingly, strains TG226 and TG227 have approximately one-half the growth rate of strains TG228, TG229, and the wildtype when *K. stuttgartiensis* encapsulin and cargo expression is induced under aerobic conditions (Figure 2). Strains TG226 and TG227 differ from strains TG228, TG229, and TG306 (wildtype) in that they encode the full-length *K. stuttgartiensis* encapsulin sequence which contains an N-terminal Sec signaling domain.

### NOC - 18 Addition Experiments: Testing for Protection from NO

In anaerobic incubations of strain TG228 (cEnc, no signalling domain) and the wild-type, there was no evidence that the *K. stuttgartiensis* encapsulin provides protection against NO. According to our hypothesis that the *K. stuttgartiensis* encapsulin would protect the cell from damage due to NO and other reactive species, strain TG228 should grow to a higher OD value post NOC-18 addition (i.e., in the presence of excess NO) than the wild-type. In opposition to our hypothesis, no difference was observed in the OD values between the wild-type and strain TG228 after NOC-18 addition (Figure 3). Instead, both strain TG228 and the wild-type TG306 plateaued after NOC-18 addition while the OD values of the unexposed samples increased with time. Experiments carried out under aerobic conditions across a broader range of NOC-18 concentrations (0.5, 1, 3, 6 mM) revealed a similar pattern (Figures S1 and S2).

**Figure 3:**
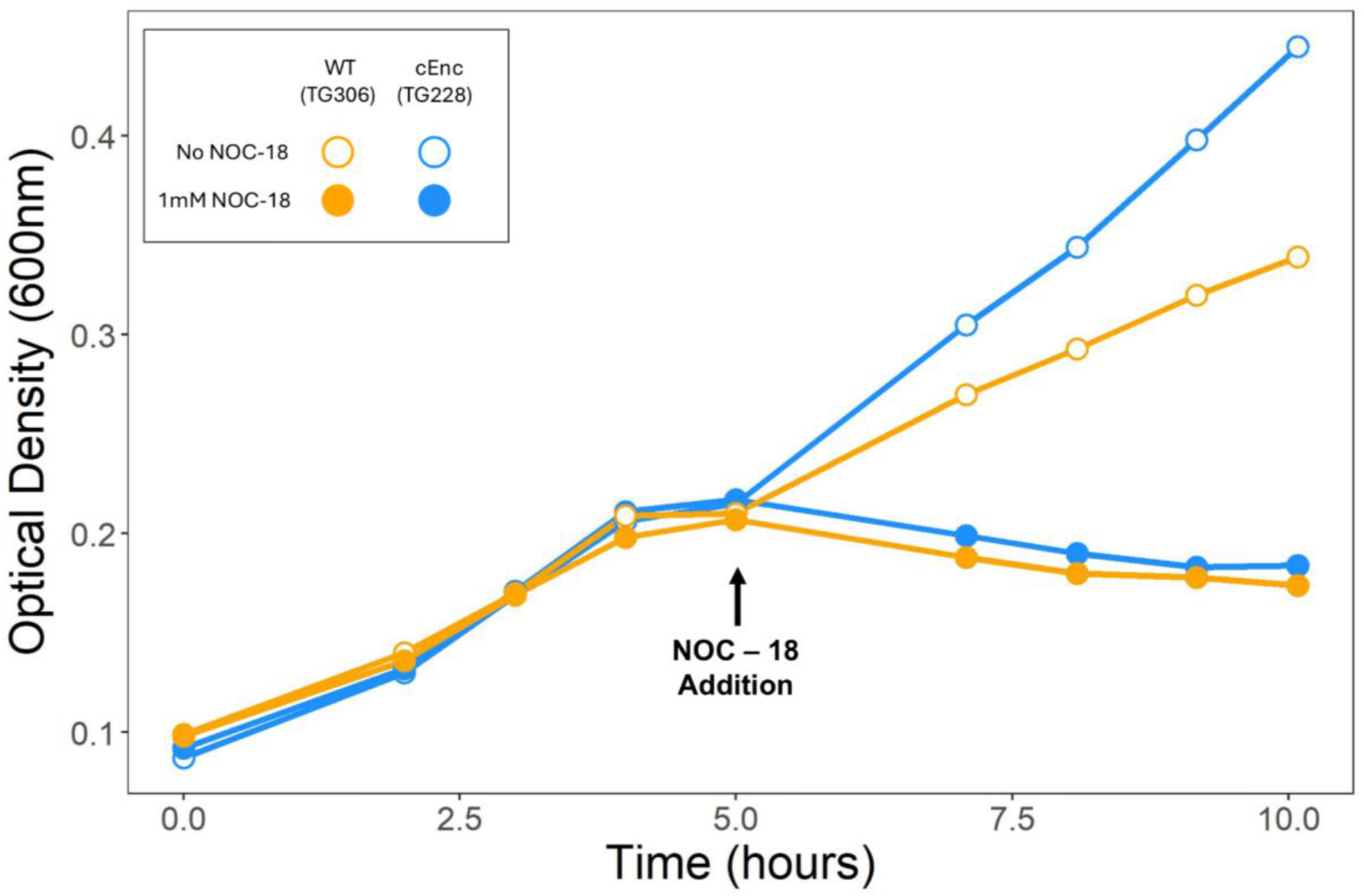
*K. stuttgartiensis* cEnc containing strain TG228 confers no protection against NO when compared to the wildtype TG306 under anaerobic conditions. Closed circles show samples when 1 mM NOC – 18 was added while open circles show samples with no added NOC – 18. NOC-18 was added at 5 hours. Strain TG306 (wildtype) is shown in orange, strain TG228 (cEnc, no signaling domain) is shown in blue. All cultures were grown at 30°C and received 2 µM hemin and FeSO_4_.

## Discussion

This study examined the biochemical functioning of anammox encapsulins using the heterologously expressed *K. stuttgartiensis* cEnc protein in an *E. coli* construct as a model system for the other anammox candidate species with similar encapsulin architectures^9,11,16^. Our experiments returned no evidence for our hypothesis that the *Kuenenia* encapsulin is an NO protection strategy, indicating that the function of the *K. stuttgartiensis* encapsulin and the other anammox encapsulins like it likely have a different, so far unidentified function.

Before directly testing our hypothesis, we performed growth experiments on a number of *E. coli* strains (Table 1) that contained combinations of both the *Kuenenia* cEnc and its putative cargo protein, a fusion NIR/HAO protein. All strains grew with high reproducibility and precision. As a result of these experiments (Figure 2) we could conclude that growth rate variability within each strain was low. An interesting result of these experiments was that strains encoding a Sec signal domain had growth rates roughly half that of strains where the Sec signal domain was removed. Sec signal domains usually transport unfolded proteins across a membrane^21^. Due to the multiple lines of evidence suggesting that anammox encapsulins are dispatched to the anammoxosome, we hypothesize that the Sec signal domain is the mechanism by which anammox encapsulins are directed there. Since *E. coli* does not have an intracellular membrane bound compartment, we hypothesize that the expressed cEnc proteins are instead exported across the inner cytoplasmic membrane. Since *E. coli* cells are not adapted to have large numbers of foreign proteins exported into the periplasm, we hypothesize that this is why Sec signal domain-containing strains have a lower growth rate than strains without the Sec signal domain. After these experiments, we chose to focus our NO protection experiments on strain TG228 which contained only the cEnc part of the putative encapsulin operon. This was because previous publications were able to show successful purification, assembly, and co-factor (heme) incorporation only for this strain. We acknowledge that testing NO protection in putative cargo producing strains is of high interest due to the putative identity of the cargo protein.

NOC - 18 addition experiments under anaerobic conditions suggest that the *Kuenenia* cEnc does not provide protection against cell damage caused by NO and possibly other reactive nitrogen species (Figure 3). These results are further supported by aerobic experiments across a broader range of concentrations (0.5, 1, 2, 6 mM NOC – 18) that also suggest that the *Kuenenia* encapsulin does not provide protection against NO (Figs. S1, S2). While our data provides valuable information that constrains the role of the *Kuenenia* encapsulin and by extension the other anammox encapsulins with similar protein sequences, we recognize possible confounding factors due to our use of heterologous proteins expressed in *E. coli*. Successful expression of cEnc, successful compartment formation and heme incorporation by cEnc were shown in a previous paper^9^; however, we do not know the degree to which all encapsulin monomers formed compartments and the degree to which heme was incorporated. In addition, heterologously expressing complex multiheme proteins in *E. coli* is not simple, especially when they contain c-type heme groups. While some successes have been reported^25,26^, studies often report challenges under both aerobic and anaerobic conditions^27^. In addition, *E. coli* possesses several native NO scavenging enzymes, including Hcp/Hcr, Hmp, and NorVW as well as the nitrite reductases NrfA and NirB which can reduce NO to NH_3_^28,29^. Based on the existing literature, NorVW and NrfA were likely active at the NOC-18 concentrations we added^29^. Complete proof that the *Kuenenia* encapsulin does not protect against NO would require deleting all these enzymes which we did not do. In summary, in the heterologous systems used here, there is no evidence for protection from NO. This does not, however, prove that NO protection does not occur in the native system. Therefore, further research is needed to identify the function of anammox encapsulin nanocompartments in both heterologous and native systems.

A promising area of future research in anammox encapsulin biochemistry would be to investigate the function of the *K. stuttgartiensis* encapsulin associated NIR/HAO. We suggest future research into this topic because anammox bacteria encode a large variety of HAO-like proteins; many of which have not been characterized. For example, *K. stuttgartiensis* encodes ten HAO-like proteins^30^, of which only two have been characterized: HDH, the HAO-like enzyme in the core anammox metabolism that oxidizes hydrazine to N_2_ gas^31^, and another unnamed *Kuenenia* HAO-like protein that generates NO from hydroxylamine (NH_2_OH)^32^. Based on experiments conducted in nitrifying organisms, it has been proposed that the presence of a C terminal tyrosine indicates whether a HAO-like protein is better suited for oxidation or reduction of nitrogenous species^33–38^. The *K. stuttgartiensis* encapsulin associated NIR/HAO lacks a C terminal tyrosine in the HAO domain^37^, indicating that this domain most likely reduces nitrogen species. A promising area of future research would be to screen for the capability to catalyze interesting reduction reactions relevant to the anammox process, such as the reduction of NO_2_^-^ to NH_4_^+^.

An additional area worthy of further experiments would be to examine if the heme containing c1 and c554 proteins located near the *K. stuttgartiensis cEnc* gene can be assembled inside the encapsulin shell as secondary cargo proteins. This hypothesis, if true, would also be able to explain the observed ∼25 nm diameter electron and iron-rich regions recorded inside the anammoxosome^18^. The *K. stuttgartiensis* genome also includes many ferritin-like proteins. Secondary cargo proteins can be located far away from the main encapsulin sequence^9^, so it is possible that one of these ferritins, if encapsulated, could explain the observed small, iron rich regions as well^39,40^.

A greater understanding of anammox biochemistry at the cellular scale would provide knowledge of the mechanisms underlying the global marine nitrogen cycle. A greater understanding of anammox biochemistry could also stimulate further technological gains that are critical for mitigating the ill effects of rising anthropogenic N inputs, which are currently estimated to be 160 Tg N yr^-1 41^ and have stimulated harmful algal blooms, induced hypoxia, and changed fisheries’ distributions and health in many coastal ecosystems^42^. Our results suggest that future attempts to understand the role of anammox encapsulins should consider hypotheses other than NO detoxification.

## Acknowledgements

We acknowledge Prof. Pam Silver for her help preparing this manuscript and throughout this project. Henry Ogilby provided valuable assistance in conducting the NOC - 18 experiments. We are grateful to the members of the Ward and Zhang labs for thoughtful comments on these experiments’ design and interpretation. We would also like to thank Prof. Xinning Zhang for the use of many pieces of equipment in her lab.

## Author contributions

T.W.G. was responsible for strain creation and proof of expression. B.B.W., J.C.T., and T.W.G. contributed to experiment design and manuscript development. J.C.T. performed all experiments and data analysis.

## Data availability statement

All raw data is publicly available at https://doi.org/10.5281/zenodo.18019779

## Additional Information

The authors report no conflicts of interest.

