## Supplementary material for "Evidence that the *Kuenenia stuttgartiensis* encapsulin does not protect against NO damage"

**Author Affiliations**

### Supplementary Materials

**Contents:**

Figure S1: Aerobic NOC-18 experiments, 0.5 and 6 mM NOC-18 additions

Figure S2: Aerobic NOC-18 experiments, 1 and 3 mM NOC-18 additions

Table S1: Number of currently known anammox encapsulins

M9 medium recipe

Protein sequence of *Kuenenia stuttgartiensis* encapsulin

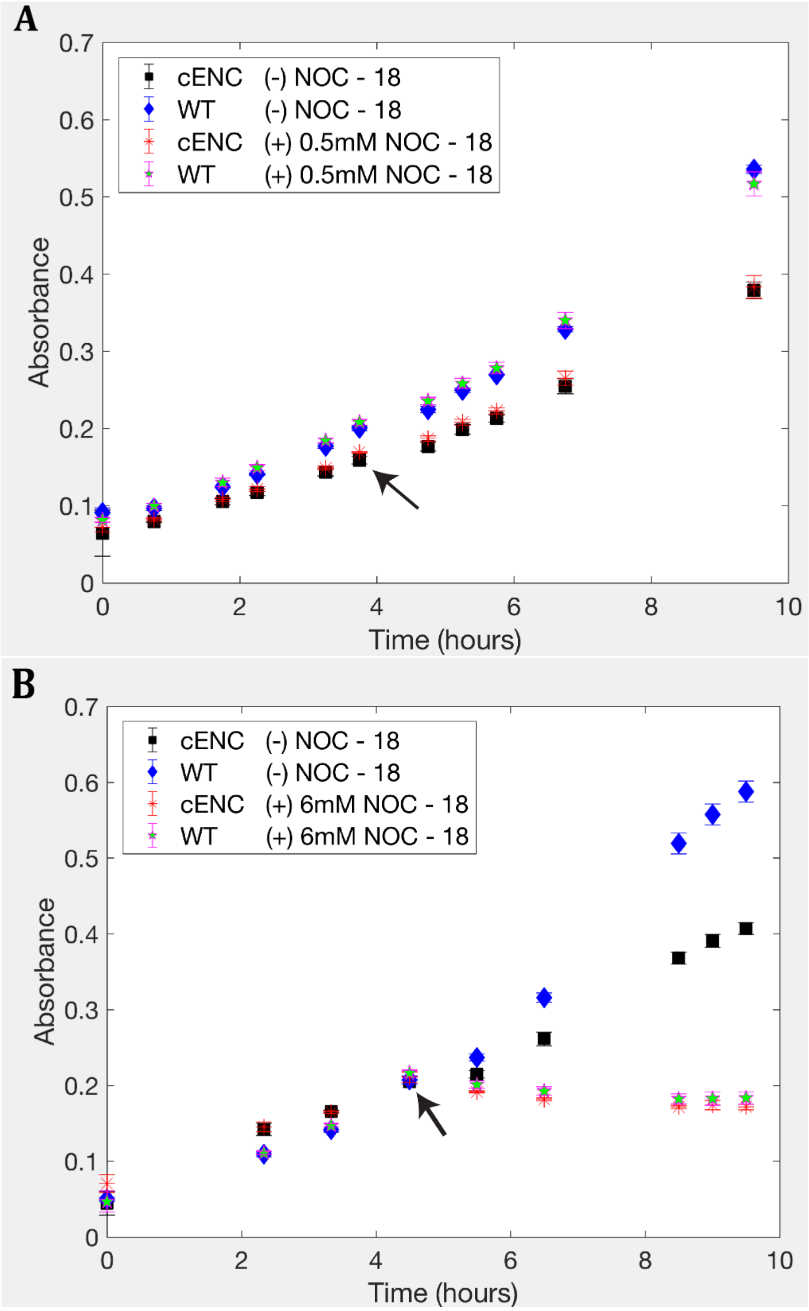


**Figure S1:** Optical density for aerobic 0.5 mM **(A)** and 6 mM **(B)** NOC - 18 addition experiments in strains TG228 (cEnc) and TG306 (WT). Arrows indicate time of NOC - 18 addition. **(A)** All cultures were grown at 30°C and received final concentrations of 2 µM hemin and 2 µM $FeSO_4$. **(B)** All cultures were grown at 37°C and received hemin and $FeSO_4$ as in Fig. S1A. Contrary to our hypothesis, at both concentrations strain TG 228 (cENC, no signaling proteins) provides no protection against NOC-18 derived NO. Points show the average optical density across triplicates at each time point and error bars show the standard deviation.

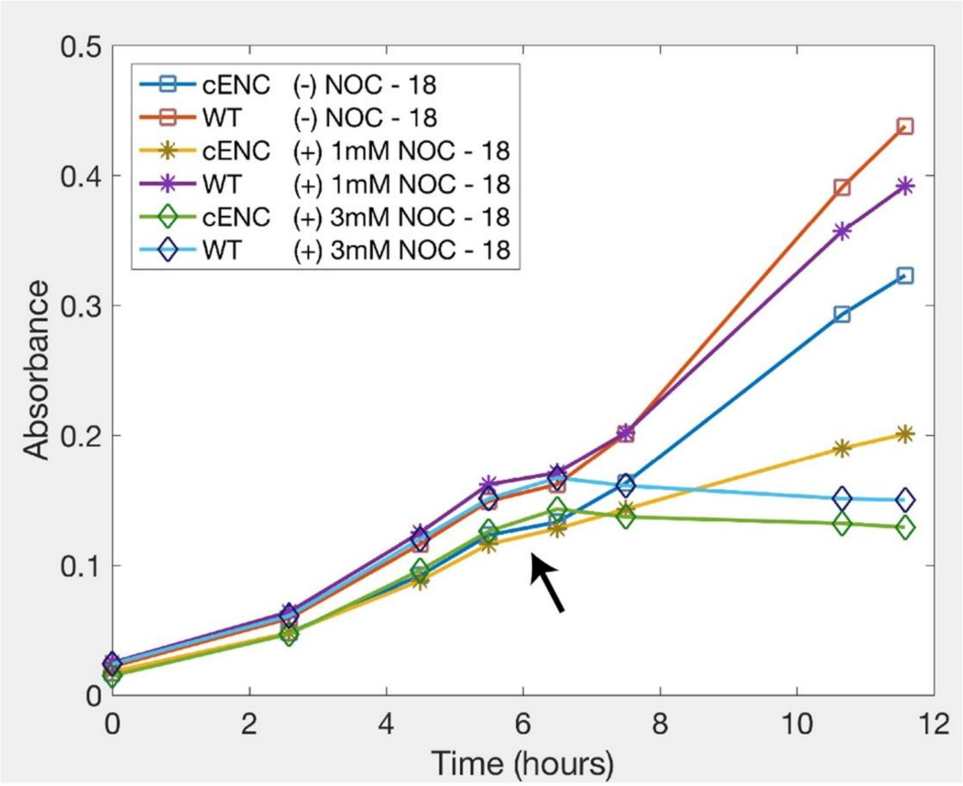


**Figure S2:** Optical density for aerobic 1 mM and 3 mM NOC - 18 addition experiments to strains TG 228 (cENC, no signaling proteins) and TG306 (WT). Arrow indicates time of NOC - 18 addition. All cultures were grown at 30°C and received final concentrations of 2 μM hemin and $FeSO_4$. Contrary to our hypothesis, at both concentrations, strain TG228 provides no protection against NOC-18 derived NO.

**Table S1: Number of currently known anammox encapsulins:**

| Anammox Species | Encapsulin Uniprot Accession | NIR/HAO fusion? | Notes |
|---|---|---|---|
| *C. Brocadia sinica JPN1*<br>*C. Brocadia sinica*<br>*C. Brocadia sp.* | A0A0C9Q3W6<br>A0A136M5G4<br>A0A399WBK9 | Yes | Encapsulin-cytochrome fusion |
| *C. Brocadia fulgida*<br>*C. Brocadia sp.* | A0A0M2UQN0<br>A0A399XHT2 | Yes | Encapsulin-cytochrome fusion |
| *C. Brocadia caroliniensis* | A0A1V4APN3 | Yes | No |
| *C. Kuenenia stuttgartiensis*<br>*C. Kuenenia stuttgartiensis* | A0A2C9CJU2<br>Q1Q6L7 | Yes | Encapsulin-cytochrome fusion |
| *C. Jettenia caeni* | I3IR75 | Yes | Encapsulin-cytochrome fusion |
| *C. Jettenia ecosi* | A0A533QDY8 | Yes | Encapsulin-cytochrome fusion |
| *C. Scalindua rubra* | A0A1E3XBX5 | No | No |
| *C. Scalindua sp. AMX11* | A0A523CPI3 | Yes | No |

The *C. Brocadia sinica JPN1, C. Brocadia sinica,* and *C. Brocadia sp.* sequences in row 2 are identical. Likewise, the *C. Brocadia fulgida,* and *C. Brocadia sp.* in row 3 are also identical. As a result, there are 8 unique anammox encapsulin protein sequences currently known.

**M9 medium recipe:**

1) Make M9 salts

   M9 Salts (5x)

   $Na_2HPO_4 \cdot 7H_2O$, 16 g
   $KH_2PO_4$, 3.75 g
   NaCl, 0.625 g
   Glycine, 5.2 g or $NH_4Cl$ 1.25g
   Deionized $H_2O$, to 0.25 liter

2) Then combine (for 500 mL)

| | |
|---|---|
| M9 salts (5X)[b] | 100 mL |
| Glucose (10%; Sigma-Aldrich)[a] | 20 mL |
| ($MgSO_4$ (1 M; Fisher Scientific)[b] | 1 mL |
| $CaCl_2$ (1 M; Fisher Scientific)[b] | 50 µL (Add last) |
| Milli-Q $H_2O$[b] | 379 mL |

[a]Filter-sterilize and store at 4°C.
[b]Autoclave before use and store at room temperature.

3) Add Ampicillin to 100 µg / mL and store at 4°C.

***Kuenenia stuttgartiensis* cEnc protein sequence:**

cEnc:

>CAJ73225.1/1363 conserved hypothetical (diheme) protein [Candidatus Kuenenia stuttgartiensis]

MVMGILNTFKKVYAVTGFFALLAVFSLSQVGSSAFAACAKVDDCFSCHTTQELNAVHK
NTPYQGQSCIVCHKAFAADDTCSDAKDGRFAKISSEININKEDWNKIQRAVHETTEKHL
VGRKFLNIYGPLGTGAQSVPLDTYGLPSWASIDMLGEGNEAIHPLKREIAQIYLIYKDFW
LFRRDIEFSKKCETPIDISAAIGAAVSVSRKEDDMVFNGLSEMGIPGLLTASGRNIMKLSD
WSVIGNGFQDVVLAVEKLTSRGFNGPFALVVSPKLYAYLHRVYERTGQLEIQGVKELV
NGGVYQSYVFNKDVALVIATGSLNMDLAVGSNYKVEYWGPQDLNHRFRVVGSSVLRI
KCPQAICTLE
Sec signal peptide is highlighted in blue
CXXCH sites are highlighted in red.
